# Cell specialization in cyanobacterial biofilm development revealed by expression of a cell-surface and extracellular matrix protein

**DOI:** 10.1101/2022.07.13.498973

**Authors:** Alona Frenkel, Eli Zecharia, Daniel Gómez-Pérez, Eleonora Sendersky, Yevgeni Yegorov, Avi Jacobs, Jennifer Benichou, York-Dieter Stierhof, Rami Parnasa, Susan S Golden, Eric Kemen, Rakefet Schwarz

## Abstract

Cyanobacterial biofilms are ubiquitous and play important roles in diverse environments, yet, understanding of the processes underlying development of these aggregates is just emerging. Here we report cell specialization in formation of *Synechococcus elongatus* PCC 7942 biofilms - a hitherto unknown characteristic of cyanobacterial multicellularity. We show that only a quarter of the cell population expresses at high levels the four-gene *ebfG*-operon that is required for biofilm formation. Almost all cells, however, are assembled in the biofilm. Detailed characterization of EbfG4 encoded by this operon revealed cell-surface localization as well as its presence in the biofilm matrix. Moreover, EbfG1-3 were shown to form amyloid structures such as fibrils and are thus likely to contribute to the matrix structure. These data suggest a beneficial ‘division of labour’ during biofilm formation where only some of the cells allocate resources to produce matrix proteins – ‘public goods’ that support robust biofilm development by the majority of the cells. Additionally, previous studies revealed the operation of a self-suppression mechanism that depends on an extracellular inhibitor, which supresses transcription of the *ebfG*-operon. Here we revealed inhibitor activity at an early growth stage and its gradual accumulation along the exponential growth phase in correlation with cell density. Data, however, do not support a threshold-like phenomenon known for quorum-sensing in heterotrophs. Together, data presented here demonstrate cell specialization and imply density-dependent regulation thereby providing novel insights into cyanobacterial communal behaviour.

**Graphical Abstract:** 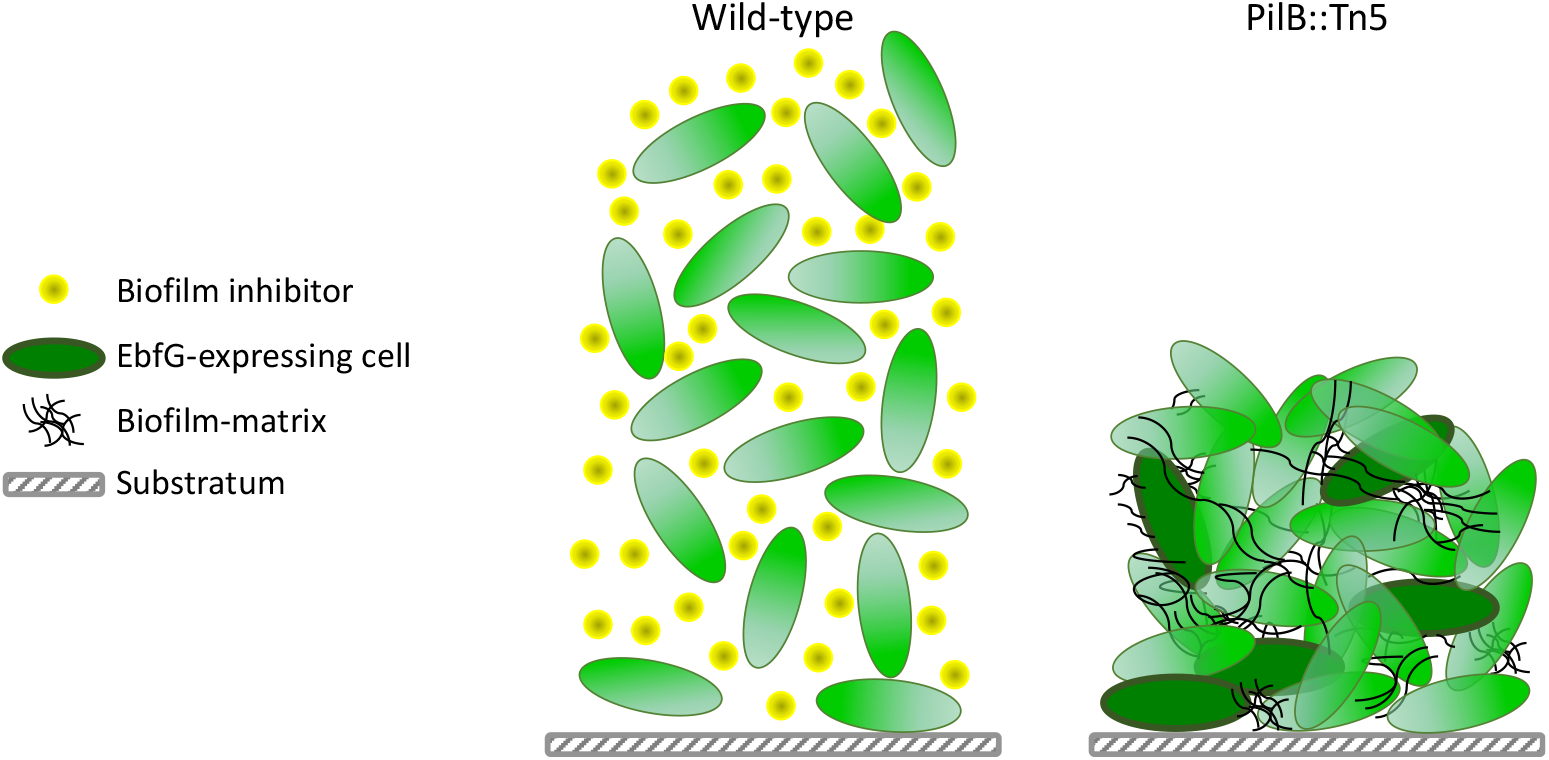

## Introduction

Cyanobacteria are highly abundant in the environment and are responsible for ∼25% of the global primary production [1, 2]. Frequently, these photosynthetic prokaryotes are found in microbial assemblages known as biofilms or part of laminated biofilms, dubbed microbial mats [3-5]. Phototrophic biofilms are often associated with industrial problems [6-8]; in contrast, such microbial consortia are beneficial e.g., for effective biomass accumulation for the biofuel industry and for harvesting of secondary metabolites [9-12]. In-depth understanding of cyanobacterial biofilm development paves the way for inhibition of deleterious biofilms and promotion of beneficial ones.

The mechanisms involved in cyanobacterial aggregation or biofilm formation started emerging only in recent years. For example, similarly to heterotrophic bacteria, cyanobacteria use the second messenger cyclic-di-GMP for regulating aggregated vs planktonic mode of growth [13]. Furthermore, the thermophilic cyanobacterium *Thermosynechococcus vulcanus* employs cyanobacteriochrome photoreceptors to mediate light-colour input for controlling cell aggregation via c-di-GMP signaling [14-16].

Microbial cells within biofilms are encased in a self-produced matrix of hydrated extracellular polymeric substances (EPS) that allows multilayering of cells and structural stability and provides a protected environment. Numerous studies in diverse heterotrophic bacteria identified particular sugar polymers and protein filaments as matrix components [17, 18], however, little is known about the cyanobacterial biofilm matrix. Yet, extracellular polysaccharides were implicated in cyanobacterial biofilm formation, for example, studies of *Synechocystis* support involvement of extracellular polysaccharides in surface adhesion [19] and cell sedimentation [20] and cellulose accumulation is responsible for cell aggregation in *T. vulcanus* RKN [21]. The exo-protein HesF of *Anabaena* sp. PCC 7120 is required for aggregation and it was proposed that it interacts with polysaccharides [22], however, its detailed role in aggregation is still unknown.

Our previous studies revealed a biofilm self-suppression mechanism in *S. elongatus* that dictates planktonic growth of this strain (Fig. 1). Inactivation of gene Synpcc7942_2071, abrogates the biofilm inhibitory process and results in a biofilm-proficient strain in contrast to the planktonic nature of WT. This gene encodes a homologue of ATPases of type 2 secretion (T2S) systems or type 4 pilus (T4P) assembly complexes, thus, the mutant was initially designated T2EΩ but recently renamed PilB::Tn5 [23-25]. The RNA chaperone Hfq and a conserved cyanobacterial protein (EbsA), which are part of the T4P complex, are also essential for the biofilm suppression mechanism [25]. Additionally, we identified four small proteins, each with a double glycine secretion motif, that enable biofilm formation (<u>e</u>nable <u>b</u>iofilm formation with a <u>G</u>G motif EbfG1-4).

**Figure 1:**
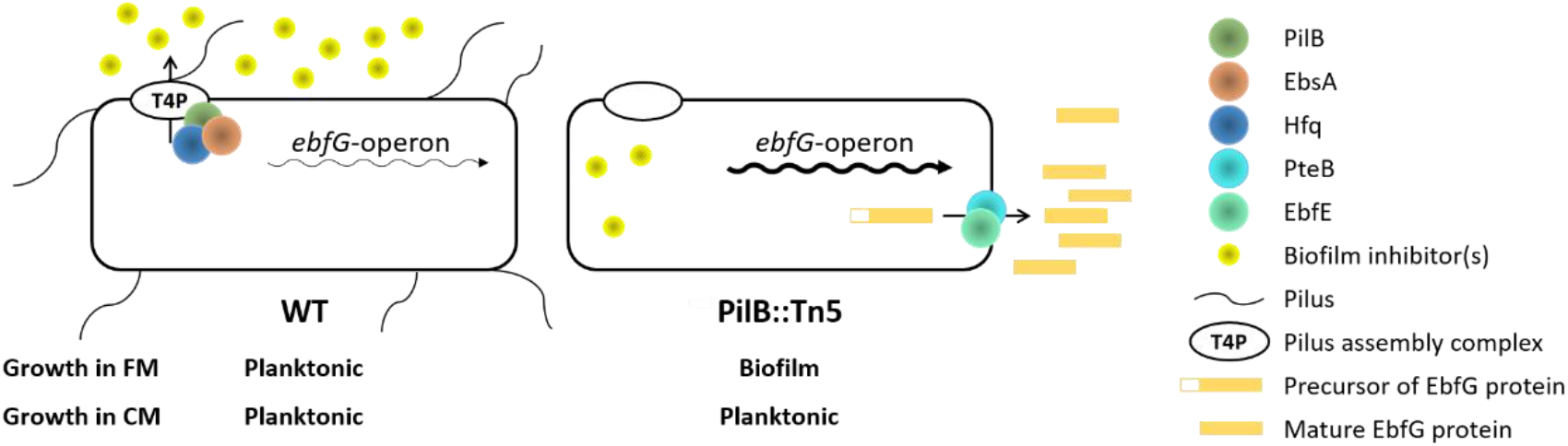
Biofilm regulation in *S. elongatus* by an extracellular inhibitor that dictates transcription of the *ebfG*-operon. The type 4 pilus (T4P) assembly complex is involved in deposition to the extracellular milieu of biofilm inhibitor(s), which dictate transcription of the *ebfG*-operon. Low and high abundance of transcripts of this operon are indicated by thin and thick arrows, respectively. FM – fresh medium; CM – conditioned medium.

The T4P complex is involved in deposition to the extracellular milieu of a biofilm inhibitor that regulates expression of the *ebfG-*operon [23, 26]. Mass-spectrometry (MS) analyses revealed the presence of EbfG1-4 in conditioned medium (CM) of PilB::Tn5 [26, 27]. Furthermore, we demonstrated that proteins PteB (<u>p</u>eptidase <u>tr</u>ansporter <u>e</u>ssential for <u>b</u>iofilm formation), which belongs to the C39 peptidases family, and EbfE (<u>e</u>nable <u>b</u>iofilm formation <u>e</u>nzyme), a homolog of microcin processing peptidases, are implicated in secretion of EbfG1-4 to the extracellular milieu [26, 28]. The role of EbfG1-4 in biofilm formation, however, was unknown. Here, using a reporter construct we demonstrate that high expression of the *ebfG*-operon is limited to a small subpopulation of cells of PilB::Tn5. Further characterization indicates cell-surface and biofilm-matrix localization of EbfG4 and strongly supports amyloid nature of EbfG2. Together, the data indicate cell specialization and imply microbial cooperation for production of extracellular components beneficial for the whole population, known as “public-goods”. Additionally, the response of the reporter strain to conditioned media harvested at different stages of logarithmic growth of the wild-type strain implies a density dependent mechanism in regulation of *S. elongatus* biofilm development.

## Materials Methods

### Strains and culture conditions and harvesting of conditioned medium (CM)

Cultures of *S. elongatus* PCC 7942 and all derived strains were grown in Pyrex tubes under bubbling with air enriched with CO_2_ as described previously [29, 30]. Construction of mutants and details of molecular manipulations are provided in Table S1. For harvesting of CM, wild-type cultures were initiated from liquid starters at OD_750_=0.2. CM harvesting was performed as described [23].

### Flow cytometry

50 ml culture at exponential phase was centrifuged (6000g, room temperature), resuspended with 4 ml fresh BG11 to obtain a concentrated culture for inoculation into fresh medium or CM at an OD_750_ of 0.5. Aliquots of 0.5 ml were taken from each culture tube following 6 days of growth and then, in case of biofilm-forming strains, planktonic cells were removed. 1.5 ml BG11 were used to resuspend the biofilmed cells by rigorous pipettation and 0.13 ml were transferred to a 1.5 ml Eppendorf tube for homogenization with a pellet pestle (Sigma-Aldrich, Z359971-1EA). The homogenized samples were filtered through a mesh (pore size 52 μm), supplemented with formaldehyde to a final concentration of 1%, diluted with PBS to OD_750_ of ∼0.0001 and measured using BD *FACSAria* (excitation 488nm, emission 530 ±30nm).

All statistical analyses were conducted in the statistical program R, version 3.3.2 [31]. FCS files obtained from *FlowJo* were analyzed with *flowcore* package [32]. Mean, median, and robust coefficient of variation (CV) of the intensity distribution for each sample were calculated. Robust CV was calculated as defined in the *FlowJo* documentation [33]. Intensity values were log-transformed. Significant difference between biofilm and planktonic cells of a particular culture was tested using Paired t-tests on several intensity distribution parameters (mean, median and robust CV). Initial analysis did not reveal significant differences between biofilm and planktonic cells within a particular culture, therefore, these data were combined for further analysis. Effect of growth medium or genetic background on intensity distribution parameters (mean, median and robust CV) was tested with 2-way repeated measures ANOVA. Specifically, mixed linear effect models were fitted with medium or genetic background as fixed effects and biological replicates as random effect, (using *lmerTest* package [34]), and the ANOVA was performed on the resulting models. Post hoc pairwise comparisons were performed by testing linear contrasts (using emmeans R package [35]), and FDR correction was applied to control for multiple testing. Normality of residuals homogeneity of variances assumptions were checked graphically.

### Dot-blot analysis

Cell extracts from 6 days old cultures were prepared as described previously [25] and 2.3 μl from diluted extracts were spotted onto TransBlot Turbo nitrocellulose membrane (Bio-Rad) and air dried for 5 min. All following procedures were performed at room temperature: Blocking was done for 1 h in 0.1% bovine serum albumin in TBST (20 mM Tris-HCl (pH 8.0) and 0.05% Tween20). Incubation with anti-FLAG (ab1162, Abcam; 1:2000 diluted in blocking solution) was performed for 1h following three washes in TBST for 5 min each. Incubation with secondary antibodies (goat anti rabbit IgG, 170-6515, Bio-Rad; 1:5000 diluted in blocking solution) was done for 1h following washes as above with extension of last wash to 15 min and signal detection using SuperSignal West Pico kit (Thermo Scientific, 34080).

### Fluorescence microscopy

Cultures were initiated and grown as described for biofilm quantification. Cultures (30 ml) were centrifuged (5 min, 6000 g, room temp) and resuspended in 1 ml phosphate-buffered saline (PBS). In case of biofilm-forming strains, the planktonic cells were removed with a pipette and the biofilmed cells were gently resuspended using 1 ml PBS. The concentrated cultures were precipitated by centrifugation in Eppendorf tubes as above, resuspended in 1 ml PBS and formaldehyde, from 16% stock solution prepared as described in Cold Spring Harbor Protocols (http://cshprotocols.cshlp.org/content/2010/1/pdb.rec12102.full), was added to a final concentration of 2%. Cells were incubated in the dark (30 min at room temperature in a tube rotator followed by 30 min incubation on ice agitation), washed once in PBS, resuspended in 1 ml PBS and the mixture was equally divided into two Eppendorf tubes. These tubes were centrifuged - cells for imaging without permeabilization were resuspended in 1 ml PBS and saved in the dark at room temperature. For imaging following permeabilization, cells were resuspended in 500 μl 0.1% triton in PBS, incubated at room temperature in a tube-rotator for 15 min and centrifuged. Cell pellet was resuspended with lysozyme solution (0.2 mg/ml dissolved in 50 mM Tris-HCl, pH 7.5 and 10 mM EDTA) and incubated for 30 min at 37°C. Cells were washed twice with 1 ml PBS. An aliquot of 200 μl was treated with an equal volume of freshly prepared blocking solution (5% BSA in PBS) in a tube-rotator for 1 h at room temperature. Cells were pelleted, resuspended in 100 μl anti-FLAG antibody (Abcam, 1:400 diluted with blocking solution), incubated for 40 min at room temperature and then 40 min at 30°C. Cells were washed twice with 100-200 μl PBS buffer, resuspended in 20 μl secondary antibody (Alexa Fluor® 488 Abcam) diluted 1:100 in blocking solution and incubated for 1h at 30°C. Pellet was washed once with 50 μl PBS and resuspended in 20 μl PBS. 3-5 μl were spread on microscopy slides prepared as follows. 10 μl of L-polylysine (Sigma) diluted 1:10 was spread on a microscope slide (approximately on a 1 cm x 1 cm region). Slides were air dried, washed by dipping them twice in double distilled water and air dried. Cells were layered on the coated area, air dried and slides were centrifuged in 50 ml falcon tubes to attach cells to the polylysine layer (300g 10 min, room temperature). 3 μl antifade [36] was spotted and covered with a coverslip. Images were recruited using Leica SP8 confocal microscope (autofluorescence: excitation - 630 nm; emission - 641 to 657 nm and detection of Alexa Fluor® 488: excitation - 488 nm; emission - 495 to 515 nm).

### Amyloid analysis

We employed the TANGO algorithm and the machine learning programs APPNN and AmyloGram for in silico amyloid prediction over the mature peptide sequence of EbfG1-4 [37-39]. The pipeline can be found at https://github.com/danielzmbp/amypred. After max-min normalization of the scores between 0 and 1, the cutoff for amyloid prediction was set at 0.5. For the annotation of amyloidogenic hotspots, we employed software with a diverse predictive background, including statistical sequence analysis (WALTZ), structural information analysis (ArchCandy and Pasta 2.0), machine-learning-based (APPNN and PATH), and metamyl, a consensus predictor [39-44]. We predicted the cross-beta three-dimensional structure from the amyloid peptide domains using Cordax and visualized it using ChimeraX [45, 46].

We used the Curli-Dependent Amyloid Generator (C-DAG) system to study amyloid formation *in vivo* [47, 48]. This system uses the built-in curli processing system from *Escherichia coli* to express recombinant proteins in order to test for their amyloid aggregation. Positive and negative controls for amyloid formation employed, the *Saccharomyces cerevisiae* prion Sup35 with aggregating domain (Sup35[NM]) and without (Sup35[M]), were encoded by pVS72 and pVS105 plasmids, respectively. EbfG proteins equipped with a CsgA secretion signal in place of the native secretion signal and fused to a 6x Histidine tag at the C-terminus were separately cloned in pExport plasmids and expressed in the *E. coli* strain VS45 (Table S1). For colony color phenotype analysis in inducing Congo Red plates (LB agar, 100 mg/l carbenicillin, 25 mg/l chloramphenicol, 0.2% w/v L-arabinose, 1mM IPTG and 10mg/l Congo Red), colonies were grown for four days at 22ºC in the dark.

In order to perform transmission electron microscopy (TEM) on the samples, we deposited cells from the inducing Congo Red plates grown for four days on copper mesh grids. After drying, we incubated with anti-Histidine primary antibody for 1 h (mouse, Sigma Aldrich) followed by incubation with secondary anti-mouse IgG conjugated to 6 nm gold beads for 1 h (goat, dianova) for the immunostained samples. All samples were negatively stained for 30 seconds in aqueous uranyl acetate, before visualization in a JSM 1400 plus transmission electron microscope (JEOL).

## Results

### ebfG-operon expression in individual cells

Previous quantitative RT-PCR analyses indicated that the *ebfG*-operon is highly transcribed in PilB::Tn5 compared to WT [26]. These data reflect the averaged transcription level, thus, to gain insight into variation within the population, here we employ a reporter construct to follow expression of this operon in individual cells. To this end, a DNA fragment bearing the putative *ebfG*-operon promoter along with the 5’ untranslated region was attached to a yellow fluorescence protein (*yfp*) gene, yielding the construct P-*ebfG*::YFP (Table S1). This fusion product was inserted in a neutral site 1 [49] in WT and PilB::Tn5 cells yielding WT/reporter and PilB::Tn5/reporter strains, respectively, which were analysed by flow cytometry (Fig. 2A&D).

**Figure 2:**
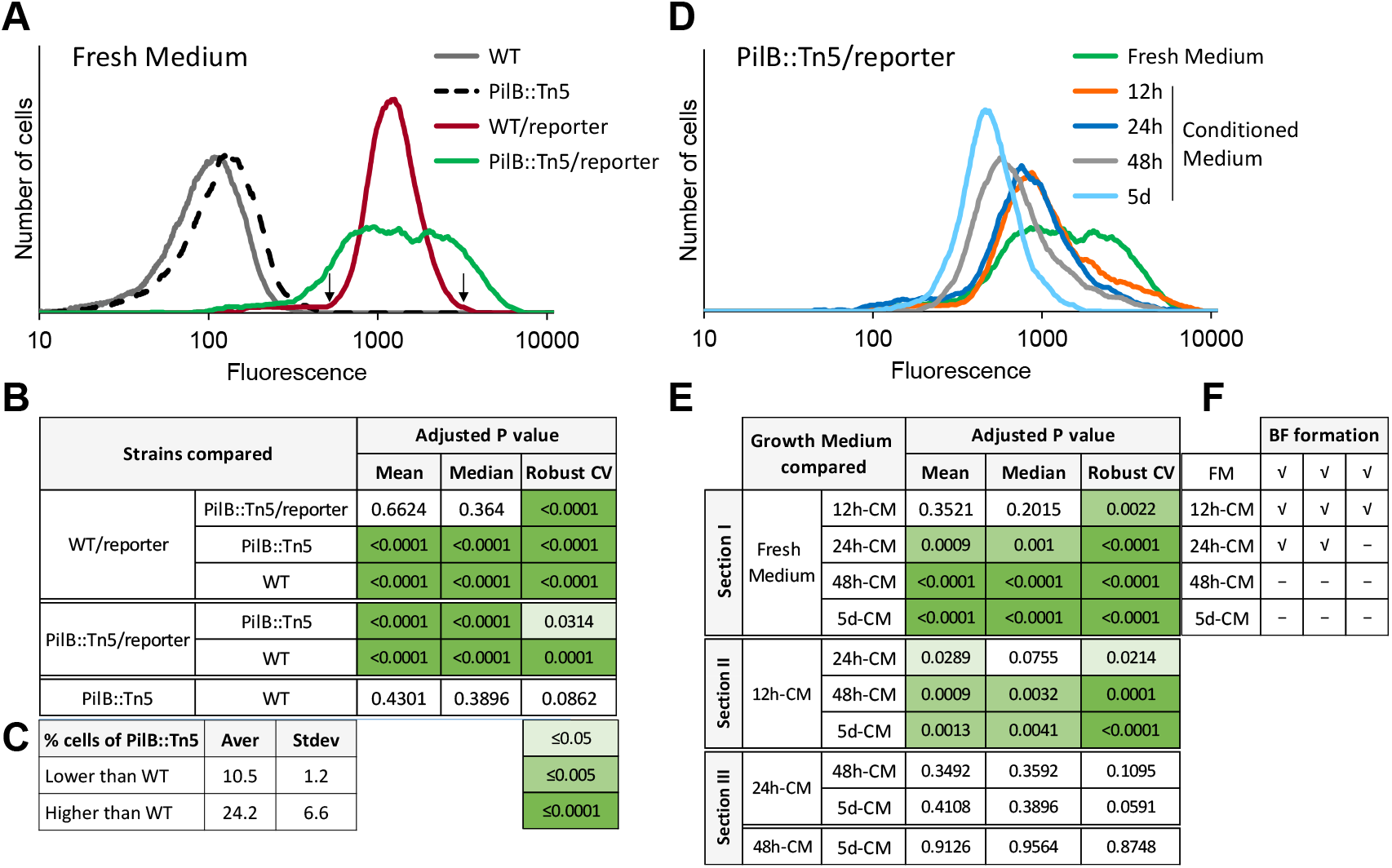
Analysis of expression of the *ebfG*-operon by flow cytometry using reporter strains. **A and B**. Number of cells as a function of fluorescence in cultures grown in fresh medium (FM) for 6 days (**A**). Strains analyzed: WT, PilB::Tn5 and their cognate reporter strains that bear a fusion of the regulatory region of the *ebfG*-operon with a yellow fluorescence protein (YFP). Arrows indicate fluorescence cutoff for calculating mutant cells with lower or higher expression of *ebfG*-operon compared to WT and summary of statistical analyses of reporter expression in FM (**B**). **C**. Fraction of PilB::Tn5/reporter cells with lower or higher expression of *ebfG*-operon compared to WT/reporter. Shown are averages and standard deviations from three independent experiments. **D and E**. Number of cells as a function of fluorescence (**D**) and summary of statistical analyses (**E**) of reporter expression in PilB::Tn5/reporter cells grown in FM and conditioned medium (CM). **B and E** show adjusted p-values of the mean, median and robust coefficient variation (rCV) from three independent experiments. **F**. Biofilm (BF) formation by PilB::Tn5/reporter cells grown in FM or CM harvested at different time points of WT culture.

Mean and median values of YFP fluorescence level is different between reporter strains and strains lacking the reporter construct, whereas these parameters are similar in WT/reporter and PilB::Tn5/reporter strains grown in fresh medium (Fig. 2B). In contrast, data dispersion is larger in PilB::Tn5/reporter compared to WT/reporter (Fig. 2A), a feature that is manifested in the significantly different robust coefficient variation (rCV, Fig. 2B). Moreover, data analysis revealed that on average, ∼10% of PilB::Tn5/reporter cells are characterized by lower - and ∼24% by higher - expression of the *ebfG*-operon (Fig. 2C). This observation suggests cell specialization in *S. elongatus* biofilm development.

Our previous studies demonstrated higher transcription of the *ebfG*-operon in PilB::Tn5 compared to WT when strains are cultured in fresh medium. Inoculation, however, of the mutant into conditioned medium (CM) from WT culture, strongly suppresses transcription [23, 26]. These data suggest involvement of intercellular communication in *S. elongatus* biofilm development possibly via a density dependent mechanism. To monitor the presence of the biofilm inhibitor along culture growth, CM was harvested from WT cultures grown for 12h, 24h, 48h and 5 days (Fig. S1) and the impact on YFP expression by PilB::Tn5/reporter was assessed. Note, conditioned media were supplemented with nutrients to negate possible nutrient limitation.

Individual flow cytometry experiments demonstrated a decrease in fluorescence intensity with CM age (e.g. Fig. 2D) in accordance with accumulation of the biofilm inhibitor during culture growth. Statistical analysis of three independent experiments indicated significant difference between rCV of cells grown in fresh medium and those grown in CM harvested at 12 h following culture initiation (12h-CM; Fig. 2E see section I of table). Additionally, mean, median and rCV were significantly different between cells grown in fresh medium and those inoculated into CM harvested at 24h, 48h and 5 days (24h-CM, 48h-CM and 5d-CM; Fig. 2E, section I of table). Moreover, mean and rCV were significantly different between cells exposed to 12h-CM and those inoculated into either 24h-CM, 48h-CM or 5d-CM (Fig. 2E, section II of table). Comparisons of the impact of CM from older cultures (Fig. 2E, section III; 24h-CM vs. 48h-CM and 5d-CM and 48h-CM vs. 5d-CM) did not reveal significant changes. Because individual experiments demonstrate accumulation of biofilm inhibitor in CM with growth time (e.g. Fig. 2D), we propose that the inhibitor is accumulated with culture age, which corresponds with culture density. Lack of significance, however, between data summarizing three biological repeats at 24h or longer culture growth, indicates variable kinetics of inhibitor accumulation in independent experiments.

Interestingly, 12h-CM had a significant impact on fluorescence rCV (Fig. 2E) and apparently, less cells expressed the *ebfG*-operon at high levels compared with fresh medium (Fig. 2D). Yet, biofilm development by these cultures (Fig. 2F) suggests that the small fraction of *ebfG*-operon highly expressing cells is sufficient to drive biofilm formation. 24h-CM significantly affected the mean, median and rCV compared with fresh medium (Fig. 2E, section I), however, in two of the three biological repeats biofilms were formed (Fig. 2F), in agreement with suggested variability in kinetics of accumulation of the biofilm inhibitor between individual experiments. 48h-CM and 5d- CM consistently inhibited biofilm formation in accordance with substantial repression of *ebfG*-operon expression under these conditions (Fig. 2E section I and 2F). Together, data indicate presence of the inhibitor at early culture stages upon initiation of the logarithmic growth (Fig. S1 and Fig. 2D and E), and suggest further accumulation with time and culture density.

### Localization of EbfG4

EbfG proteins do not share primary sequence similarity or domains with proteins of known function. To get insight into the role of these proteins in biofilm development, we selected EbfG4 for further characterization. This particular EbfG protein was chosen because a mutational approach impairing the secretion motif of either one of the EbfG proteins revealed that the mutation in EbfG4 had the most prominent impact on biofilm development compared with EbfG1-3 [26]. To follow EbfG4 localization in biofilm-forming and planktonic strains we introduced a FLAG-epitope tagged EbfG4 (EbfG4::FLAG) to the double mutant having inactivations in both *pilB* and Synpcc7942_1134 (PilB::Tn5/EbfG4Ω). The double mutant grows planktonically – similarly to WT, 100% of the chlorophyll is in planktonic cells as assessed by measurement of the relative amount of chlorophyll in suspension of total chlorophyll in the culture (Fig. 3). Insertion of a DNA fragment bearing either the native *ebfG*-operon or one encoding EbfG4::FLAG into the double mutant (PilB::Tn5/EbfG4Ω/Comp and PilB::Tn5/EbfG4Ω/EbfG4::FLAG, respectively), restored biofilm development; similarly to PilB::Tn5, less than 5% of the chlorophyll is in suspended cells (Fig. 3). This analysis validated functionality of the tagged EbfG4.

**Figure 3:**
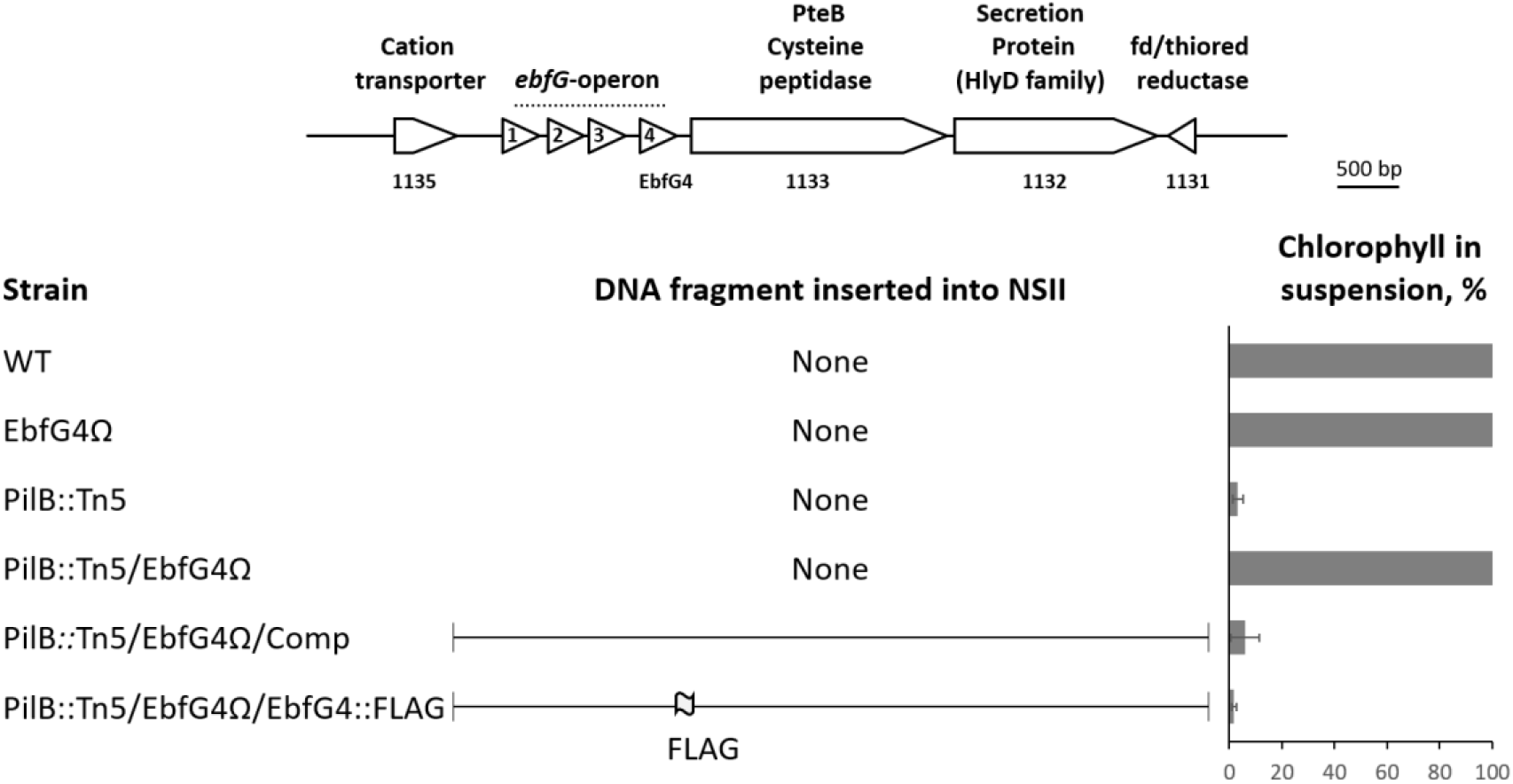
FLAG-tagged EbfG4 is functional in biofilm development. Genomic region of the *ebfG*-operon. Bar graph presents percentage of total chlorophyll in the suspended cells (average of three independent biological repeats ± standard deviation). Strains analyzed include: WT, a strain in which *ebfG*4 was insertionally inactivated (EbfG4Ω), PilB::Tn5, the double mutant PilB::Tn5/EbfG4Ω, double mutants complemented with the indicated fragments encoding native or FLAG-tagged EbfG4 (PilB::Tn5/EbfG4Ω/Comp and PilB::Tn5/EbfG4Ω/EbfG4::FLAG, respectively.

A first indication that EbfG4 is localized to the cell surface was obtained by dot-blot analysis. Whole cells and cell extracts were spotted onto a nitrocellulose-membrane and probed with anti-FLAG antibodies (Fig. 4). This analysis suggested association of EbfG4 with the cell surface, as revealed by the signal obtained from whole cells (Fig. 4, PilB::Tn5/EbfG4Ω/EbfG4::FLAG). In contrast, the ATPase of T4P complex known to be localized cytoplasmically, was not detected in whole cells but only in cell extracts (Fig. 4, PilB::Tn5/PilB::FLAG), thus confirming availability of internal epitopes for detection only in cell extracts. EbfG4 was not detected in whole cells or in cell extracts of the planktonic strain EbfG4Ω/EbfG4::FLAG (Fig. 4), thus, EbfG4 is neither secreted nor accumulated internally when the biofilm suppression mechanism is active.

**Figure 4:**
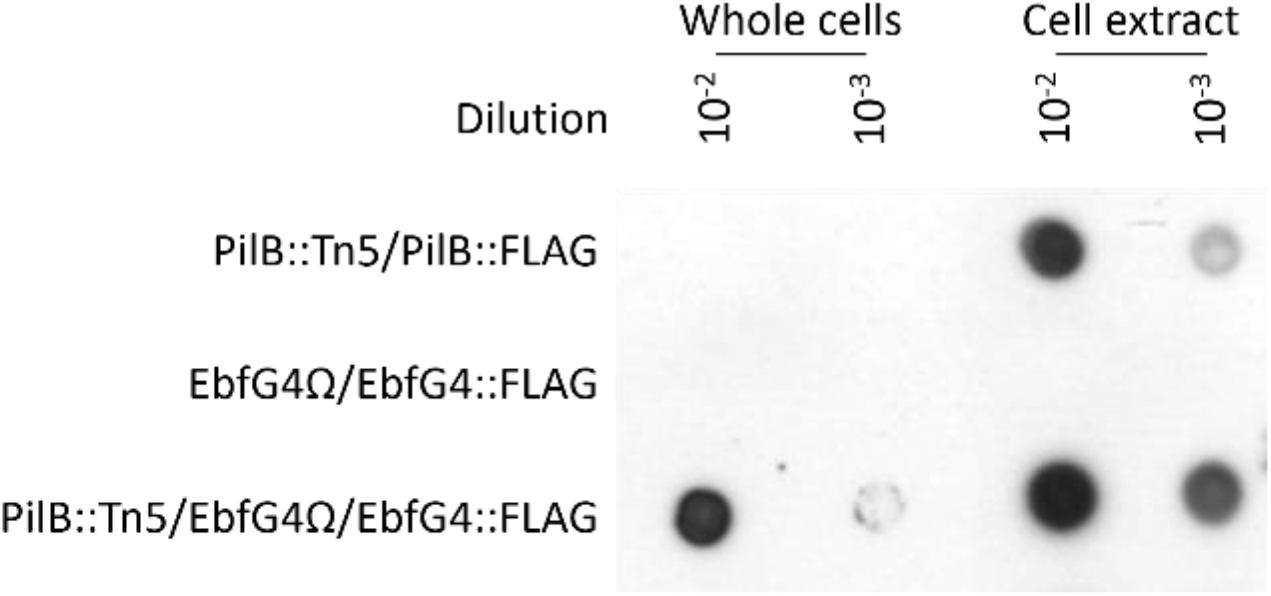
Dot-blot analysis using anti-FLAG antibodies. Whole cells and cell extracts of the following strains were analyzed: pilB-inactivated strain complemented with FLAG-tagged PilB (PilB::Tn5/PilB::FLAG), ebfG4-inactivated strain complemented with FLAG-tagged EbfG4 (EbfG4Ω/EbfG4::FLAG) and the latter strain that also harbors pilB inactivation (PilB::Tn5/EbfG4Ω/EbfG4::FLAG). Strains PilB::Tn5/PilB::FLAG and EbfG4Ω/EbfG4::FLAG are planktonic and PilB::Tn5/EbfG4Ω/EbfG4::FLAG forms biofilm.

To follow up on the observation suggesting cell surface association of EbfG4 (Fig. 4), we examined by immunocytochemistry non-permeabilized cells that were subjected to anti-FLAG antibodies, thus allowing detection of only extracellular EbfG4. Green signal representing EbfG4 was detected only in strain PilB::Tn5/EbfG4Ω/EbfG4::FLAG but was absent from the cognate control strain PilB::Tn5/EbfG4Ω/Comp confirming specific detection of the FLAG epitope (Fig. 5). Moreover, the green signal is associated with clustered cells (Fig. 5, middle and right columns), while unclustered cells lack green signal and are visualized only by autofluorescence (Fig. 5).

**Figure 5:**
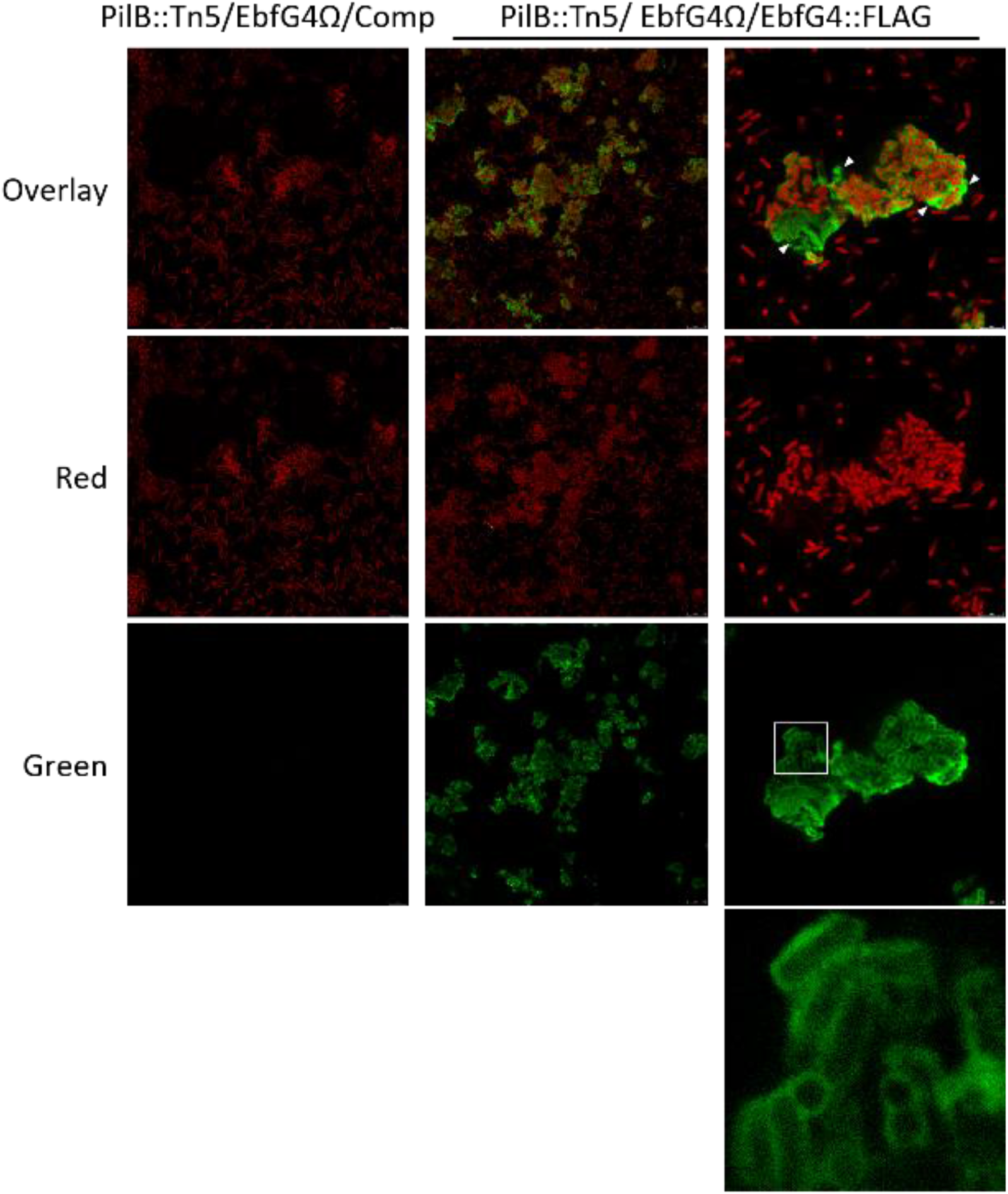
Visualization of EbfG4::FLAG in non-permeabilized cells by immunocytochemistry of confocal fluorescence microscopy. Strains analyzed: PilB::Tn5/EbfG4Ω/Comp and PilB::Tn5/EbfG4Ω/EbfG4::FLAG. Red (excitation 630nm; emission 636-695nm) represents autofluorescence whereas green (excitation 490nm; emission: 497-539nm) indicates presence of EbfG4::FLAG. White square indicates the enlarged area shown in the panel below. Arrowheads point at patches of green fluorescence in areas void of cells.

Closer examination revealed green signal encasing many of the clustered cells indicating EbfG4 localization throughout the cell surface (Fig. 5, right column). Additionally, patches of green fluorescence are observed in areas void of cells (Fig. 5, right column, arrowheads), and 3D-imaging clearly indicates the presence of EbfG4 in between cells (Fig. 6) supporting its role as a matrix protein.

**Figure 6:**
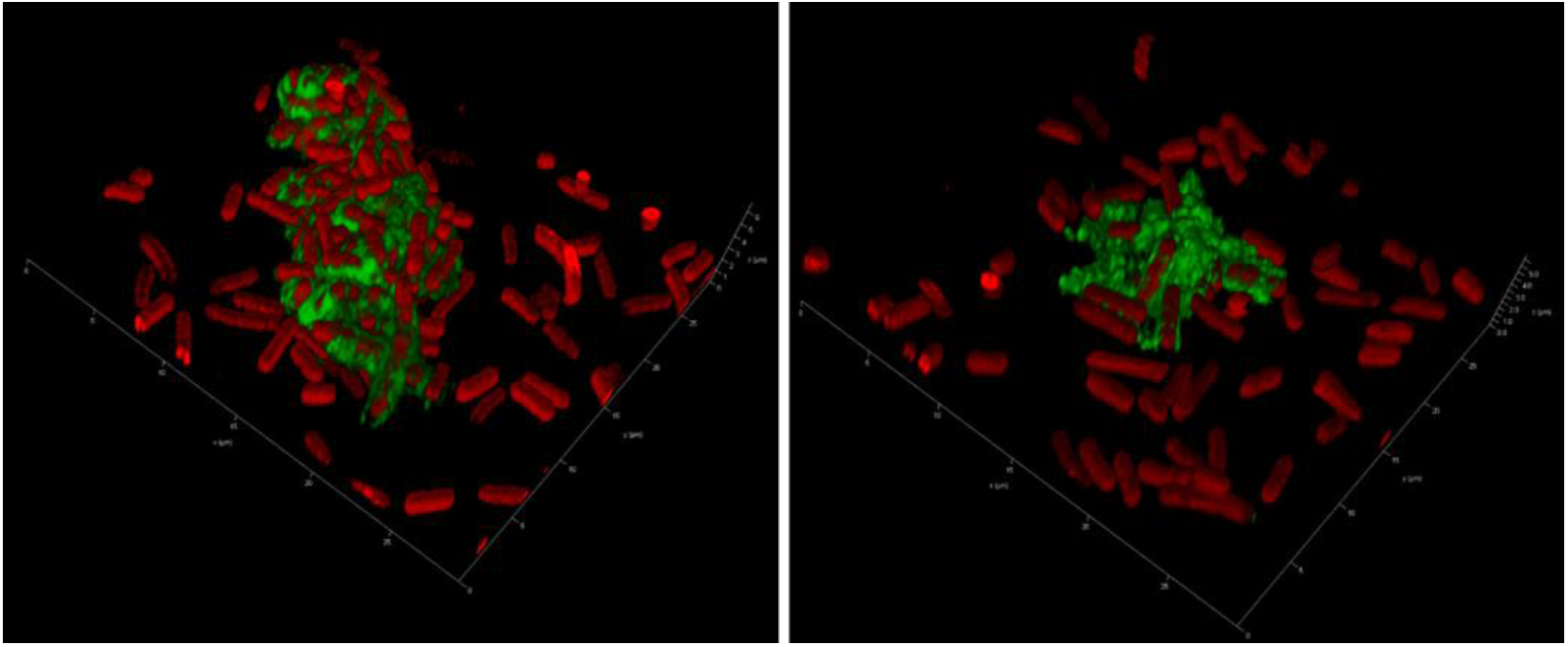
3D-imaging of strain PilB::Tn5/EbfG4Ω/EbfG4::FLAG. See details in legend to Fig. 5.

Next, we visualized permeabilized cells to test whether EbfG4 is also detected intracellularly. Images similar to those observed without permeabilization emerged from these analyses (Fig. 7). Close examination of an area mostly void of extracellular green fluorescence did not reveal green signal within the cells (Fig. 7, right column).

**Figure 7:**
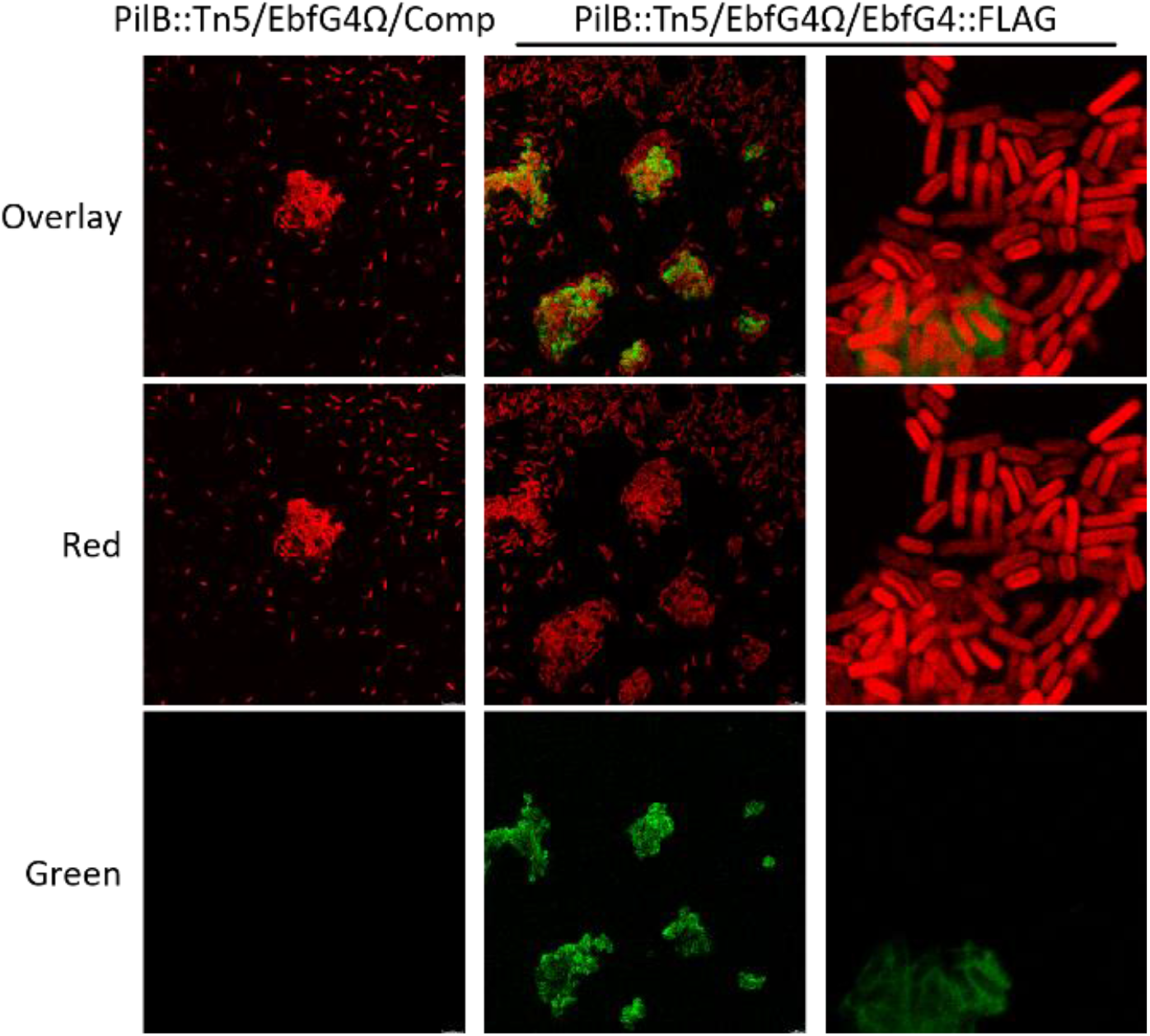
Visualization of EbfG4::FLAG in permeabilized cells by immunocytochemistry and confocal fluorescence microscopy. Strains analyzed: PilB::Tn5/EbfG4Ω/Comp and PilB::Tn5/EbfG4Ω/EbfG4::FLAG. Red (excitation 630nm; emission 636-695nm) represents autofluorescence whereas green (excitation 490nm; emission: 497-539nm) indicates presence of EbfG4::FLAG.

Successful visualization of internal PilB::FLAG, exclusively in permeabilized cells, validate penetration of the antibodies used for detection (Fig. S2). Together, these data indicate that EbfG4 does not accumulate intracellularly to detectable levels and suggest efficient secretion of this protein.

### Examination of amyloid formation by products of the ebfG-operon

To test whether EbfG proteins might contribute to biofilm matrix formation through insoluble aggregates, we initially performed an *in silico* characterization of their tendency to fold as amyloids using a pipeline that consists of several prediction software to build consensus models. We found that EbfG2 and 3 had the highest prediction score, which was above the 0.5 cut-off for classification into amyloid-forming proteins. Using separate tools with diverse prediction methods for the identification of amyloid hotspots within the amino acid sequence, we found that the highly predicted regions occurred at the same location in the alignment of EbfG2 and EbfG3. Despite variations in the individual sequences, the core hotspot followed the motif AΦNIΠ, where Φ represents a hydrophobic residue, F or I, and Π, a small side chain residue, G or S (Fig. 8A). Such motif also occurs in EbfG1, however, in EbfG4 a polar uncharged amino acid (glutamine) is present instead of a hydrophobic residue (AQNIG). We modelled the amyloidogenic LAFNIG peptide from EbfG2 for its ability to form cross-beta structures. We found that the hexapeptide arranged in a steric zipper of antiparallel fashion (Figure S3), characteristic of amyloid proteins [50].

**Figure 8:**
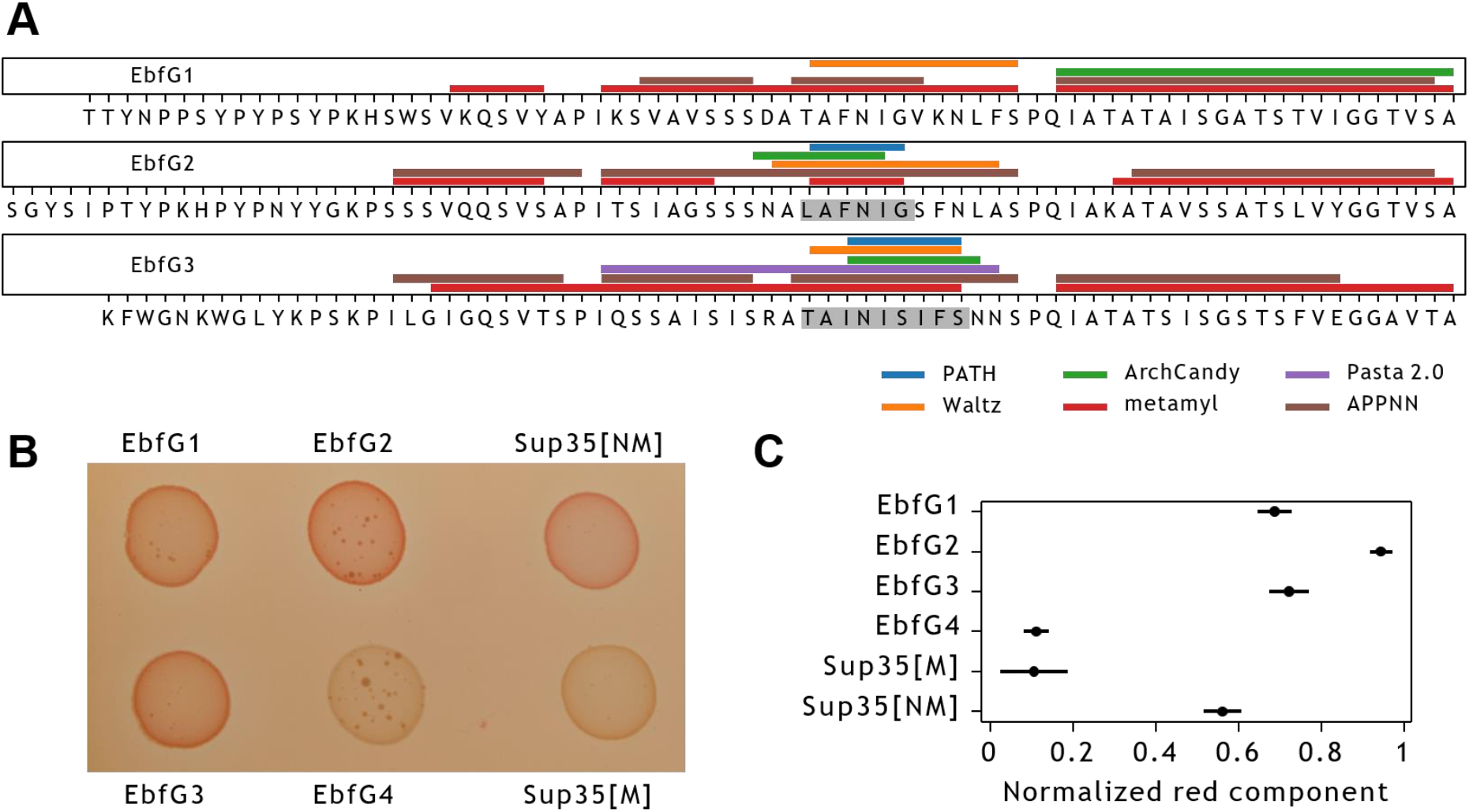
Amyloid prediction and heterologous expression of EbfG1-4 proteins. **(A)** Prediction of amyloidogenic hotspots in the sequences of EbfG1-3 using six software prediction tools. Shaded fragments correspond to consensus positive prediction in at least four of the predictors. **(B)** Colony color phenotype of EbfG-expressing *Escherichia coli* compared to negative (Sup35[M]) and positive (Sup35[M]) controls for amyloid formation in inducing plates. (C) Normalized red component of the brim of each colony as analyzed in ImageJ, based on five biological replicates and displayed as the mean and 95% confidence interval (t distribution).

To support these predictions, we heterologously expressed the EbfG proteins to test for the formation of amyloids *in vivo*. To this end, we cloned the mature proteins upstream of the GG motif in the Curli-Dependent Amyloid Generator (C-DAG) system fused to a 6x Histidine Tag at the C-terminus (Table S1). After induction with arabinose, we found a phenotype for amyloid formation in EbfG1, 2 and 3, evident by the colour shift of the colonies due to the binding of Congo Red, similarly to the positive control (Fig. 8B). Consistent with the *in silico* predictions, EbfG2 showed the strongest phenotype while EbfG4 showed no sign of amyloid formation, comparable to the negative control staining (Fig. 8C).

When looking at the induced EbfG2-producing C-DAG cells under the transmission electron microscope, we detected fibril-like structures in the extracellular space. The fibrils resemble those produced by the positive control, however shorter in length and associated with vesicle-like structures (Fig. 9A). Using immunostaining methods, we corroborated the identity of the fibrils as containing EbfG2 protein, based on the colocalization of gold nano-beads directed to the 6x Histidine tag from the protein. We observed unevenness in the distribution of the labelling which could be due to variable antibody accessibility of the tag, both for the positive control and EbfG2 (Fig. 9B).

**Figure 9:**
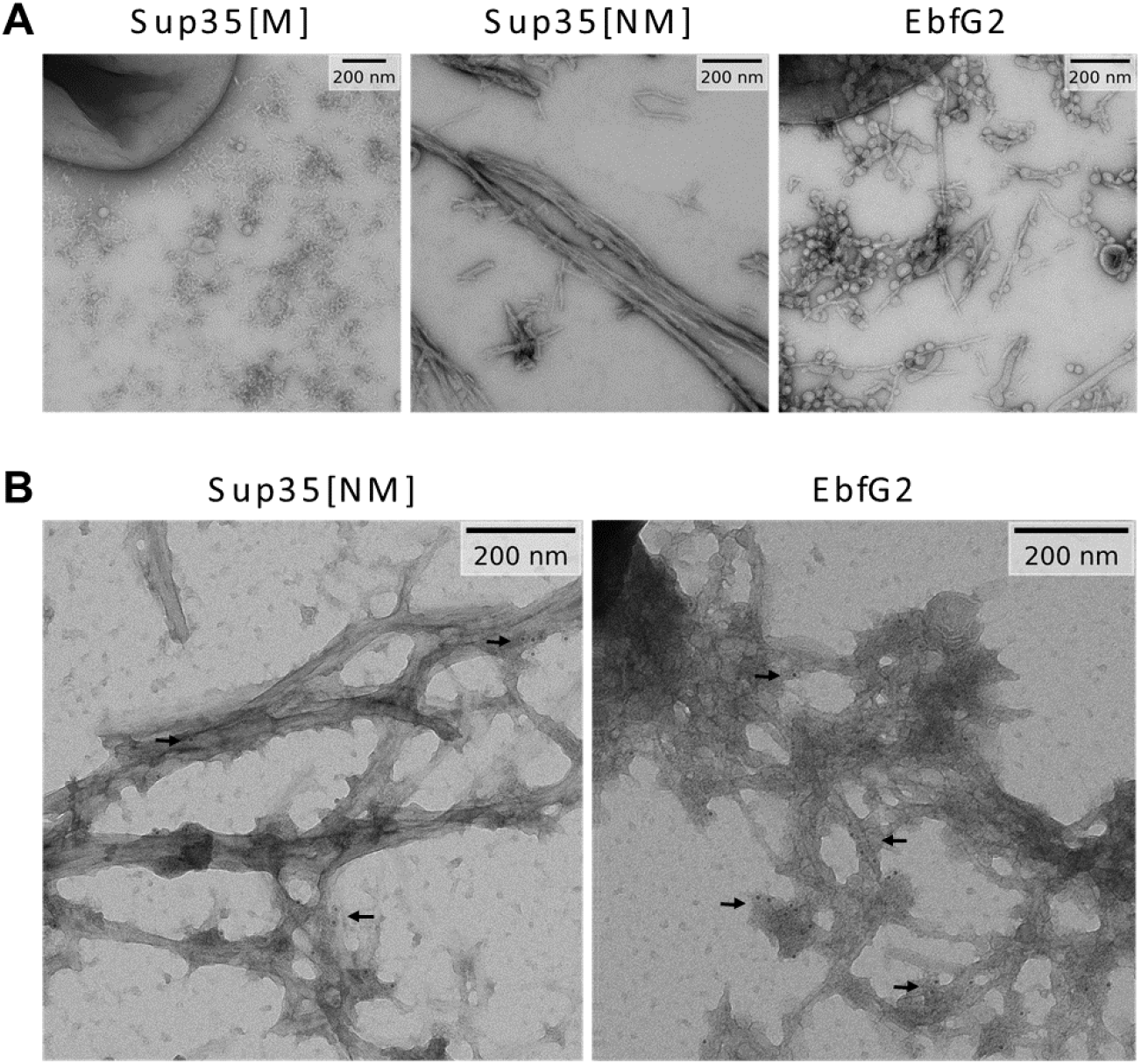
Transmission electron microscopy images of EbfG2 expressed in the C-DAG system. **(A)** Fibril-like structures are found in the extracellular space for the positive control (Sup35[NM]) and EbfG2 but not for the negative control (Sup35[M]) when expressed in the C-DAG system. **(B)** Immunostained samples where 6 nm gold particles localize to the fibril-like structures of the positive control and EbfG2.

## Discussion

### Cell specialization and density-dependent regulation of public goods production in S. elongatus biofilm development

Previous genetic and point mutation analyses demonstrated the requirement of the EbfG proteins for biofilm formation [23, 26]. As manifested by analysis in individual cells, expression of the operon encoding these genes varied substantially more in the biofilm-forming mutant PilB::Tn5 than among WT cells (Fig. 2A-D). Although ∼75% of the mutant cells express the *ebfG*-operon at similar or lower level compared to WT (Fig. 2C), ∼90% of the mutant cells are found in the biofilm (Fig. 3, [23, 26, 28]). These findings are consistent with cell specialization during *S. elongatus* biofilm development. Given that EbfG1-3 are prone to form amyloids (Fig. 8) and TEM analysis revealed that EbfG2 forms fibrillar structures similar to amyloids, we propose that a relatively small subpopulation of the mutant culture produces matrix components that support robust biofilm development by the majority of the cells. Taken together, we propose cell specialization in production of so-called public goods. Such a phenomenon may confer selective advantage because only a minority of the cells invest resources for the benefit of the population, which gains protection within the biofilm [51]. Cell specialization in microbes have been documented in heterotrophic bacteria, for example, during matrix formation in *Bacillus subtilis* [52]. Such differentiation, however, was not previously reported for cyanobacterial biofilm development, thus, this study, which uncovers cell differentiation suggests division of labor in communities of these ecologically important photosynthetic prokaryotes.

Comparisons of expression of the *ebfG*-operon in fresh medium and under CM harvested at different time points along growth of WT culture indicate production and secretion of the inhibitor at early culture stages and suggest further accumulation with time and cell density. The amount of biofilm inhibitor present following 12h growth is sufficient to affect reporter expression (Fig. 2D and E), yet is insufficient for biofilm inhibition (Fig. 2F). Up to 24h the inhibitor gradually accumulates (Fig. 2E) and at 48h the inhibitor reaches levels that consistently inhibit biofilm development (Fig. 2F). Together, data imply a density-dependent mechanism, however, a gradual impact of the inhibitor is observed rather than a “threshold-like” effect typical of quorum-sensing mechanisms known in heterotrophic bacteria, e.g. induction of the lux-operon [53-56]. Cyanobacterial quorum-sensing is largely unknown, although N-octanoyl homoserine lactone was suggested to be involved in quorum-sensing in *Gloeothece* PCC6909 [57]. Biosynthetic LuxI-like proteins, which are responsible for production of acylated homoserine lactones in numerous heterotrophs, however, are not encoded in the majority of cyanobacterial genomes [58], therefore, such molecules are not likely to represent a general mechanism for cyanobacterial intercellular communication. An additional study demonstrated a governing role of extracellular signals produced at high density on transcription of particular genes in low-density cultures of *N. punctiforme* PCC 73102. These data support the existence of a quorum-sensing-like mechanism(s), however, the nature of the signal(s) and the regulatory network have yet to be identified [58].

### Dual function of EbfG4 – cell surface and matrix protein

Microscopic analysis revealed that some of the cells of PilB::Tn5/EbfG4Ω/EbfG4::FLAG present the EbfG4 protein on their cell surface (Figs. 5&7). Interestingly, EbfG4 is only observed on the surface of clustered cells and cells that lack EbfG4 labelling are dissociated from the clusters (Figs. 5&7). Together, these observations are in accordance with an adhisin function of EbfG4. Additionally, EbfG4 was observed in the intercellular space (Figs. 5-7), consistent with the hypothesis that it serves as a matrix component. It is possible that similarly to the adhisin SasG of *Staphylococcus aureus* [59], EbfG4 is initially deposited to the cell surface and later is shed to the extracellular matrix. The role of EbfG4 in the matrix is unknown, however, although it does not form amyloids by itself (Fig. 8) it may be associated with amyloid structures formed by EbfG1-3.

### Amyloids as part of S. elongatus biofilm matrix

When establishing biofilms, microbes require a resilient scaffold on which the cells can settle. Bacteria from diverse ecosystems have solved this problem by producing and releasing functional amyloids into their environment [60, 61]. Amyloid proteins are able to assemble into long and strong fibrils, which can withstand chemical and physical stresses [62]. However, the production of amyloids is a process that can easily get out of control, therefore, it requires a complex and dedicated machinery for appropriate manufacturing. Here, we have investigated the amyloid forming capabilities of the *ebfG*-operon proteins and found strong evidence supporting amyloid formation in EbfG1-3, and most manifestly in EbfG2. EbfG4, which has a prominent role in biofilm formation, however, did not spontaneously form amyloid fibrils. Consistent with homology in the amyloid hotspots, we hypothesize this could be related to a mechanism intended to control aggregation. By separating the amyloid nucleators, in this case EbfG1-3, from other components of the fibril, e.g. EbfG4, better control over the synthesis of amyloids could be achieved. This would be analogous to, for example, the functioning of CsgB and CsgA in the production of curli, the biofilm backbone in *E. coli* [63]. As observed in the TEM pictures, the EbfG2 fibrils were much shorter than the positive control and did not bundle together. Heterogeneous fibrils formed of several EbfG proteins could result in more stable fibrils, as seeding of amyloids composed of perfect repeats has been shown to cause fragmentation [64].

Given the amyloid nature of EbfG1-3 proteins one may speculate that these matrix components of *S. elongatus* biofilms assist in recruitment of other microbes for establishment of multispecies biofilms. Additionally, our findings pave the way for controlling formation of unwanted biomats, by using amyloid disrupting compounds, as already shown for other bacteria [65, 66]. On the other hand, intentional use of protein seeds could facilitate stronger amyloids and hence elicit formation of beneficial biofilms.

## Supporting information

EbfG_Supplemental data

## Acknowledgments

We thank Ryan Simkovsky for providing the vector for EbfG4-tagging. Studies in the laboratories of Rakefet Schwarz and Susan Golden were supported by the program of the National Science Foundation and the US-Israel Binational Science Foundation (NSF-BSF 2012823). This study was also supported by grants from the Israel Science Foundation (ISF 1406/14 and 2494/19) to Rakefet Schwarz. Studies in the laboratory of Eric Kemen were supported by the graduate school GRK 1708 “Molecular principles of bacterial survival strategies” and the European Research Council (ERC) under the DeCoCt research program (grant agreement: ERC-2018-COG 820124).

## References

1. Falkowski, P.G., The role of phytoplankton photosynthesis in global biogeochemical cycles. Photosynth. Res., 1994. 39(3): p. 235–258.

2. Flombaum, P., et al., Present and future global distributions of the marine Cyanobacteria Prochlorococcus and Synechococcus. Proc Natl Acad Sci U S A, 2013. 110(24): p. 9824–9.

3. Bolhuis, H., M.S. Cretoiu, and L.J. Stal, Molecular ecology of microbial mats. FEMS Microbiol Ecol, 2014. 90(2): p. 335–50.

4. Rossi, F. and R. De Philippis, Role of cyanobacterial exopolysaccharides in phototrophic biofilms and in complex microbial mats. Life (Basel), 2015. 5(2): p. 1218–38.

5. Veach, A.M. and N.A. Griffiths, Testing the light:nutrient hypothesis: Insights into biofilm structure and function using metatranscriptomics. Molecular Ecology, 2018. 27(14): p. 2909–2912.

6. Wagner, M. and A. Loy, Bacterial community composition and function in sewage treatment systems. Curr. Opin. Biotechnol., 2002. 13(3): p. 218–27.

7. Ivnitsky, H., et al., Bacterial community composition and structure of biofilms developing on nanofiltration membranes applied to wastewater treatment. Water Res., 2007. 41(17): p. 3924–3935.

8. Belila, A., et al., Bacterial community structure and variation in a full-scale seawater desalination plant for drinking water production. Water Res, 2016. 94: p. 62–72.

9. Heimann, K., Novel approaches to microalgal and cyanobacterial cultivation for bioenergy and biofuel production. Curr Opin Biotechnol, 2016. 38: p. 183–189.

10. Strieth, D., R. Ulber, and K. Muffler, Application of phototrophic biofilms: from fundamentals to processes. Bioprocess Biosyst Eng, 2018. 41(3): p. 295–312.

11. Bruno, L., et al., Characterization of biofilm-forming cyanobacteria for biomass and lipid production. J Appl Microbiol, 2012. 113(5): p. 1052–64.

12. Egan, S., T. Thomas, and S. Kjelleberg, Unlocking the diversity and biotechnological potential of marine surface associated microbial communities. Curr. Opin. Microbiol., 2008. 11(3): p. 219–25.

13. Agostoni, M., C.M. Waters, and B.L. Montgomery, Regulation of biofilm formation and cellular buoyancy through modulating intracellular cyclic di-GMP levels in engineered cyanobacteria. Biotechnol Bioeng, 2016. 113(2): p. 311–319.

14. Enomoto, G., et al., Cyanobacteriochrome SesA Is a Diguanylate Cyclase That Induces Cell Aggregation in Thermosynechococcus. Journal of Biological Chemistry, 2014. 289(36): p. 24801–24809.

15. Enomoto, G., et al., Three cyanobacteriochromes work together to form a light color-sensitive input system for c-di-GMP signaling of cell aggregation. Proceedings of the National Academy of Sciences of the United States of America, 2015. 112(26): p. 8082–8087.

16. Enomoto, G., Y. Okuda, and M. Ikeuchi, Tlr1612 is the major repressor of cell aggregation in the light-color-dependent c-di-GMP signaling network of Thermosynechococcus vulcanus. Sci Rep, 2018. 8(1): p. 5338.

17. Branda, S.S., et al., Biofilms: the matrix revisited. Trends Microbiol., 2005. 13(1): p. 20–6.

18. Flemming, H.C. and J. Wingender, The biofilm matrix. Nature Reviews Microbiology, 2010. 8(9): p. 623–633.

19. Fisher, M.L., et al., Export of extracellular polysaccharides modulates adherence of the Cyanobacterium Synechocystis. PLoS One, 2013. 8(9): p. e74514.

20. Jittawuttipoka, T., et al., Multidisciplinary evidences that Synechocystis PCC6803 exopolysaccharides operate in cell sedimentation and protection against salt and metal stresses. PLoS One, 2013. 8(2): p. e55564.

21. Kawano, Y., et al., Cellulose accumulation and a cellulose synthase gene are responsible for cell aggregation in the cyanobacterium Thermosynechococcus vulcanus RKN. Plant Cell Physiol, 2011. 52(6): p. 957–66.

22. Oliveira, P., et al., HesF, an exoprotein required for filament adhesion and aggregation in Anabaena sp PCC 7120. Environmental Microbiology, 2015. 17(5): p. 1631–1648.

23. Schatz, D., et al., Self-suppression of biofilm formation in the cyanobacterium Synechococcus elongatus. Environ Microbiol, 2013. 15(6): p. 1786–94.

24. Nagar, E. and R. Schwarz, To be or not to be planktonic? Self-inhibition of biofilm development. Environmental Microbiology, 2015. 17(5): p. 1477–1486.

25. Yegorov, Y., et al., A Cyanobacterial Component Required for Pilus Biogenesis Affects the Exoproteome. mBio, 2021. 12(2).

26. Parnasa, R., et al., Small secreted proteins enable biofilm development in the cyanobacterium Synechococcus elongatus. Sci Rep, 2016. 6: p. 32209.

27. Nagar, E., et al., Type 4 pili are dispensable for biofilm development in the cyanobacterium Synechococcus elongatus. Environ Microbiol, 2017. 19(7): p. 2862–2872.

28. Parnasa, R., et al., A microcin processing peptidase-like protein of the cyanobacterium Synechococcus elongatus is essential for secretion of biofilm-promoting proteins. Environ Microbiol Rep, 2019. 11(3): p. 456–463.

29. Sendersky, E., et al., Quantification of Chlorophyll as a Proxy for Biofilm Formation in the Cyanobacterium Synechococcus elongatus. Bio-protocol, 2017. 7(14): p. https://www.bio-protocol.org/e2406 xDOI:10.21769/BioProtoc.2406.

30. Suban, S., et al., Impairment of a cyanobacterial glycosyltransferase that modifies a pilin results in biofilm development. Environ Microbiol Rep, 2022. 14(2): p. 218–229.

31. Team, R.C. R: A language and environment for statistical computing. https://www.R-project.org/ 2021.

32. Ellis B H.P., Hahne F, Le Meur N, Gopalakrishnan N, Spidlen J, Jiang M, Finak G flowCore: Basic structures for flow cytometry data. R package version 2.0.1. 2020.

33. https://docs.flowjo.com/flowjo/workspaces-and-samples/ws-statistics/ws-statdefinitions/.

34. Kuznetsova, A., Per B. Brockhoff, and Rune HB Christensen, lmerTest package: tests in linear mixed effects models. Journal of statistical software 2017. 82(13): p. 1–26 R package version 3.1-0.

35. Lenth, R.V. emmeans: Estimated Marginal Means, aka Least-Squares Means.. R package version 1.6.2-1 2021.

36. Johnson, G.D. and G.M. Nogueira Araujo, A simple method of reducing the fading of immunofluorescence during microscopy. J Immunol Methods, 1981. 43(3): p. 349–50.

37. Fernandez-Escamilla, A.M., et al., Prediction of sequence-dependent and mutational effects on the aggregation of peptides and proteins. Nat Biotechnol, 2004. 22(10): p. 1302–6.

38. Burdukiewicz, M., et al., Amyloidogenic motifs revealed by n-gram analysis. Sci Rep, 2017. 7(1): p. 12961.

39. Familia, C., et al., Prediction of Peptide and Protein Propensity for Amyloid Formation. PLoS One, 2015. 10(8): p. e0134679.

40. Oliveberg, M., Waltz, an exciting new move in amyloid prediction. Nat Methods, 2010. 7(3): p. 187–8.

41. Ahmed, A.B., et al., A structure-based approach to predict predisposition to amyloidosis. Alzheimers Dement, 2015. 11(6): p. 681–90.

42. Walsh, I., et al., PASTA 2.0: an improved server for protein aggregation prediction. Nucleic Acids Res, 2014. 42(Web Server issue): p. W301–7.

43. Wojciechowski, J.W. and M. Kotulska, PATH - Prediction of Amyloidogenicity by Threading and Machine Learning. Sci Rep, 2020. 10(1): p. 7721.

44. Emily, M., A. Talvas, and C. Delamarche, MetAmyl: a METa-predictor for AMYLoid proteins. PLoS One, 2013. 8(11): p. e79722.

45. Louros, N., et al., Structure-based machine-guided mapping of amyloid sequence space reveals uncharted sequence clusters with higher solubilities. Nat Commun, 2020. 11(1): p. 3314.

46. Pettersen, E.F., et al., UCSF ChimeraX: Structure visualization for researchers, educators, and developers. Protein Sci, 2021. 30(1): p. 70–82.

47. Sivanathan, V. and A. Hochschild, A bacterial export system for generating extracellular amyloid aggregates. Nat Protoc, 2013. 8(7): p. 1381–90.

48. Sivanathan, V. and A. Hochschild, Generating extracellular amyloid aggregates using E. coli cells. Genes & Development, 2012. 26(23): p. 2659–2667.

49. Taton, A., et al., Broad-host-range vector system for synthetic biology and biotechnology in cyanobacteria. Nucleic Acids Res, 2014. 42(17): p. e136.

50. Sawaya, M.R., et al., Atomic structures of amyloid cross-beta spines reveal varied steric zippers. Nature, 2007. 447(7143): p. 453–7.

51. Yin, W., et al., Biofilms: The Microbial “Protective Clothing” in Extreme Environments. International Journal of Molecular Sciences, 2019. 20(14).

52. Dragos, A., et al., Division of Labor during Biofilm Matrix Production. Curr Biol, 2018. 28(12): p. 1903–1913 e5.

53. Fuqua, C. and E.P. Greenberg, Listening in on bacteria: acyl-homoserine lactone signalling. Nat. Rev. Mol. Cell Biol., 2002. 3(9): p. 685–695.

54. Parsek, M.R. and E. Greenberg, Sociomicrobiology: the connections between quorum sensing and biofilms. Trends Microbiol., 2005. 13(1): p. 27–33.

55. Kolter, R. and E.P. Greenberg, Microbial sciences - The superficial life of microbes. Nature, 2006. 441(7091): p. 300–302.

56. Bassler, B.L., Small talk. Cell-to-cell communication in bacteria. Cell, 2002. 109(4): p. 421–424.

57. Sharif, D.I., et al., Quorum sensing in Cyanobacteria: N-octanoyl-homoserine lactone release and response, by the epilithic colonial cyanobacterium Gloeothece PCC6909. ISME J, 2008. 2(12): p. 1171–82.

58. Guljamow, A., et al., High-Density Cultivation of Terrestrial Nostoc Strains Leads to Reprogramming of Secondary Metabolome. Appl Environ Microbiol, 2017. 83(23).

59. Geoghegan, J.A., et al., Role of surface protein SasG in biofilm formation by Staphylococcus aureus. J Bacteriol, 2010. 192(21): p. 5663–73.

60. Gomez-Perez, D., et al., Amyloid Proteins in Plant-Associated Microbial Communities. Microb Physiol, 2021. 31(2): p. 88–98.

61. Taglialegna, A., I. Lasa, and J. Valle, Amyloid Structures as Biofilm Matrix Scaffolds. Journal of Bacteriology, 2016. 198(19): p. 2579–2588.

62. Rambaran, R.N. and L.C. Serpell, Amyloid fibrils: abnormal protein assembly. Prion, 2008. 2(3): p. 112–7.

63. Hammer, N.D., J.C. Schmidt, and M.R. Chapman, The curli nucleator protein, CsgB, contains an amyloidogenic domain that directs CsgA polymerization. Proceedings of the National Academy of Sciences, 2007. 104(30): p. 12494–12499.

64. Rasmussen, C.B., et al., Imperfect repeats in the functional amyloid protein FapC reduce the tendency to fragment during fibrillation. Protein Sci, 2019. 28(3): p. 633–642.

65. Romero, D., et al., Biofilm Inhibitors that Target Amyloid Proteins. Chemistry & Biology, 2013. 20(1): p. 102–110.

66. Jain, N., et al., Inhibition of curli assembly and Escherichia coli biofilm formation by the human systemic amyloid precursor transthyretin. Proc Natl Acad Sci U S A, 2017. 114(46): p. 12184–12189.

