## Supplementary material for "Cell specialization in cyanobacterial biofilm development revealed by expression of a cell-surface and extracellular matrix protein": EbfG_Supplemental data

#### Supplementary Information

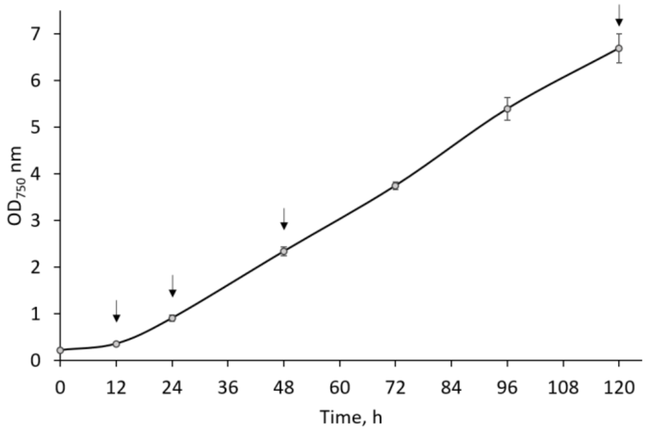


**Figure S1: Growth of WT cultures as measured by $OD_{750}$ nm as a function of time.** Data represent averages and standard deviations from 3 technical repeats. Arrows indicate time points at which CM was harvested.

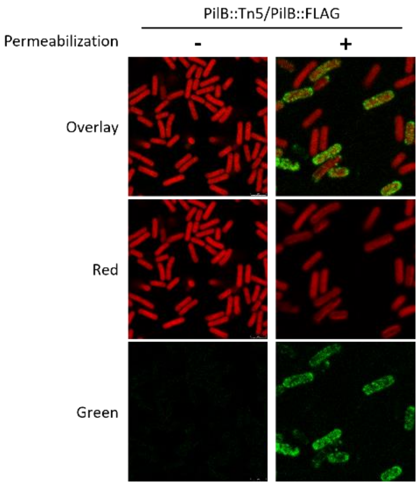


**Figure S2: Immunocytochemistry with or without permeabilization of strain PilB::Tn5/PilB::FLAG.**

Red (excitation 630nm; emission 636-695nm) represents autofluorescence whereas green (excitation 490nm; emission: 497-539nm) indicates presence of EbfG4::FLAG.

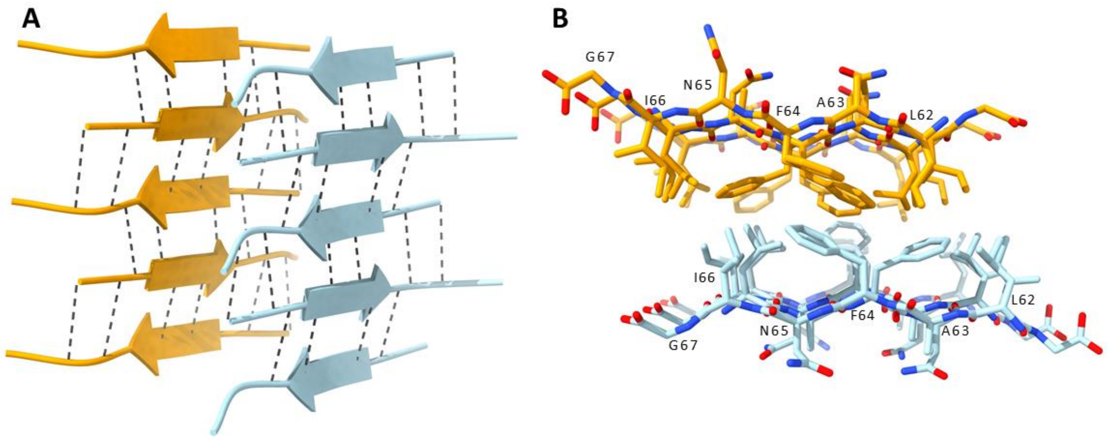


**Figure S3: Cross-beta structure modelling of amyloidogenic LAFNIG peptide from EbfG2.**

(A) Three-dimensional model of the LAFNIG peptide displaying a cross-beta antiparallel structure. In dashed black lines, hydrogen bonds holding together each cross-beta sheet are depicted. (B) Upper view of the structure highlighting the steric zipper, where hydrophobic residues (L, A, F and I) are concentrated in between the beta-sheets and hydrophilic residues (N and G) directed towards the outside.

| Cloning of FLAG-tagged EbfG4* | | |
|---|---|---|
| **Primer Name** | **Primer sequence<br>(upper forward, lower reverse)** | **Purpose** |
| RP10-swaI-F | CGGCCAATAACCCAGGGATTTTGTA<br>CTGCGACTCGACCAAG | *Cloning FLAG Fragment #1* |
| RP10-swaI-R | CTGCCGGGGAGCTCCTTCATTTGTT<br>ATGAACGTTGGCGATCG | *Cloning Fragment #3* |
| EbfG4-FLAG-R | CTTATCGTCGTCATCCTTGTAATCAA<br>GATTGACAGCTACGTTAATGG | *Cloning FLAG Fragment #1* |
| EbfG4-FLAG-F | GATTACAAGGATGACGACGATAAGT<br>AGCCAGTGCGATCGCAAG | *Cloning FLAG Fragment #2* |
| NS2-2F | CTCTTGGTGCTGTTCAGTC | *Sequencing* |

| Cloning of YFP-reporter ** | | |
|---|---|---|
| **Primer name** | **Primer sequence<br>(upper forward, lower reverse)** | **Purpose** |
| P-EbfG::YFP Forw | AGTCGGCCAATAACCCAGGGATTT<br>TGTACTGCGACTCGACCAAG | P-EbfG::YFP, *Fragment #1* |
| P-EbfG_RBS::YFP tail Rev | CACCATCTTAAGACCTCCTTTATTT<br>TTTTTGATTCGCCCTTGACTC | |
| P-EbfG:: YFP_Forw | AGTCGGCCAATAACCCAGGGATTT<br>TGTACTGCGACTCGACCAAG | P-EbfG::YFP, *Fragment #2* |
| YFP_genomic 3' UTR Rev | CATCCGTCAGGATGGCCTTCTCCT<br>GCAGGGCCGAGTTTGTACAAGAA<br>AGC | |
| NSI- pRP22 ins F | ACCGGTTAATAGTACCTGTG | P-EbfG::YFP, PCR analysis on *E. coli* cells |
| NSI- pRP22 ins R | ATGCCCGCGACATCTTCC | |
| NSI and pRP22 F | AGACGAGGCAAGCATTGAGC | P-EbfG::YFP, PCR analysis on cyano cells |
| NSI- pRP22 ins R | ATGCCCGCGACATCTTCC | |

| Cloning of fragments encoding EbfG proteins into C-DAG system# | | |
|---|---|---|
| **Primer name** | **Primer sequence<br>(upper forward, lower reverse)** | **Purpose** |
| pExport:ebfg1 | TATAGCGGCCGCAAAATTCTGGGG<br>G | PCR amplification |
| | TATATCTAGATTAGTGATGATGGTG<br>ATGGTGGGCAGATACAGTTC | |
| pExport:ebfg2 | TATAGCGGCCGCAAGTGGTTATTCC<br>ATT | PCR amplification |
| | TATATCTAGATTAGTGATGATGGTG<br>ATGGTGAGCTGAAACAGTC | |
| pExport:ebfg3 | TATAGCGGCCGCAAAATTCTGGGG<br>G | PCR amplification |
| | TATATCTAGATTAGTGATGATGGTG<br>ATGGTGGGCTGTAACTGC | |
| pExport:ebfg4 | TATAGCGGCCGCAACTAATTGCAAT<br>CC | PCR amplification |
| | TATATCTAGATTAGTGATGATGGTG<br>ATGGTGTTGCGATTATATTTCTT | |
| pExport | CCTGACGCTTTTTATCG | Sequencing |
| | GCTGAAAATCTTCTCTC | |

#### Table S1: Summary of cloning information

* EbfG4::FLAG tagged vectors were generated using the GeneArt® Seamless Cloning and Assembly Kit (Life Technologies) and Top10 cells (Life Technologies) on three PCR-generated fragments per plasmid and a SwaI-digested CYANO-VECTOR cloning plasmid, pCV0049 or pAM4937, that encodes for kanamycin (Km) resistance, Neutral Site II (NS2) integration, and

a *ccdB*-suicide gene that is removed upon SwaI digestion [1]. PCR fragments were amplified from the complementation plasmid pAM4997 (pRP10, [2]) using Q5® High-Fidelity DNA polymerase (New England Biolabs) and primers designed to add overlaps for cloning into the SwaI cut site or to add the flag tags to the C-terminus of the encoded EbfG4 protein. Clones were screened by PCR and the entire complementation region of the newly generated vectors were verified by Sanger sequencing.

** The genomic region bearing the promoter of the *ebfG*-opreon and YFP were PCR amplified and then served as primers for each other to PCR amplify a fusion product.

### PCR products were amplified from pRP10 [2] and cloned into the vector pExport [3, 4]. Fragments were inserted at the NotI (5') and XbaI (3') sites between the CsgA secretion signal and stop codon of pExport. All cloning products were validated by PCR analyses and sequencing.

#### References


1. Taton, A., et al., *Broad-host-range vector system for synthetic biology and biotechnology in cyanobacteria.* Nucleic Acids Res, 2014. **42**(17): p. e136.
2. Parnasa, R., et al., *Small secreted proteins enable biofilm development in the cyanobacterium Synechococcus elongatus.* Sci Rep, 2016. **6**: p. 32209.
3. Sivanathan, V. and A. Hochschild, *A bacterial export system for generating extracellular amyloid aggregates.* Nat Protoc, 2013. **8**(7): p. 1381-90.
4. Sivanathan, V. and A. Hochschild, *Generating extracellular amyloid aggregates using E. coli cells.* Genes & Development, 2012. **26**(23): p. 2659-2667.
